# Antisense oligonucleotides targeting Valosin-containing protein improve muscle pathology and molecular defects in cell and mouse models of multisystem proteinopathy

**DOI:** 10.1101/2025.07.25.665261

**Authors:** Pallabi Pal, Michele Carrer, Lan Weiss, Olga G. Jaime, Cheng Cheng, Alyaa Shmara, Victoria Boock, Danae Bosch, Marwan Youssef, Yasamin Fazeli, Megan Afetian, Tamar R. Grossman, Michael R. Hicks, Paymaan Jafar-nejad, Virginia Kimonis

## Abstract

Valosin-containing protein (VCP) related disease, also known as multisystem proteinopathy 1 (MSP1), is an autosomal dominant disease caused by gain-of-function pathogenic variants of the *VCP* gene. The disease is associated with inclusion body myopathy, early-onset Paget’s disease of bone, frontotemporal dementia, and familial amyotrophic lateral sclerosis. There is currently no treatment for this progressive disease associated with early demise resulting from proximal limb girdle and respiratory muscle weakness. We hypothesize that regulating VCP hyperactivity to normal levels can reduce the disease pathology. In this study, we assessed the effect of antisense oligonucleotides (ASOs) specifically targeting the human *VCP* gene in the patient (R155H) iPSC-derived skeletal muscle progenitor cells (SMPCs). ASOs were well tolerated up to a concentration of 5 µM, and significantly reduced VCP protein expression in the SMPCs by 48% (95% CI [39–56]). We also treated the transgenic mouse model of VCP disease with the overexpressed humanized VCP gene severe A232E pathogenic variant with weekly subcutaneous ASO injections starting from 6 months of age for 3 months. In the skeletal muscle of transgenic mice, ASOs resulted in 30% (95%CI [27–32]) knockdown of VCP protein compared to control ASO. The ASO-mediated reduction of VCP expression in muscle tissue was associated with improvement in autophagy flux and reduction in TDP-43 expression, hallmarks of VCP disease. In addition, ASO-treated VCP A232E mice showed improvements in functional tests of muscle strength, such as rotarod and inverted screen test compared to mice treated with control ASO. These results suggest that targeting VCP could be beneficial in preventing the progression of the VCP myopathy and hold promise for the treatment of patients with VCP MSP1.

## Introduction

Multisystem proteinopathy 1 (MSP1) or inclusion body myopathy associated with Paget’s disease of the bone, frontotemporal dementia (IBMPFD), and amyotrophic lateral sclerosis (ALS) is a rare syndromic disease caused by gain-of-function variants in the Valosin Containing Protein (*VCP*) gene [1, 2]. Over 85 pathogenic variants have been identified in more than 400 patients worldwide, 80–90% of whom have myopathy predominantly affecting the proximal girdle muscle, the diaphragm, and other muscle groups [3]. Paget disease of bone is present in 49% of patients and affects the pelvis, skull, scapulae, and vertebral column, resulting in pain, bone deformities, and fractures [4, 5]. Additionally, 27% of patients have premature frontotemporal dementia and manifest signs of neurodegeneration in the frontal and temporal lobes, including dysnomia, comprehension deficits, and social unawareness, and approximately 15% of patients have ALS [6, 7]. There are currently no effective treatments available for the neuromuscular components of MSP1, and patients generally die from respiratory failure and cardiomyopathy, typically in their 50s to 60s. Therefore, development of a treatment for this devastatingly progressive VCP-related neuromuscular disease is urgently needed.

VCP is a ubiquitously expressed hexameric protein belonging to the ATPases associated with various activities (AAA+) chaperone-like protein family. VCP is a critically important component of the ubiquitin-proteasome system. Mutation in one VCP allele can lead to a wide array of dysfunctions, including improper protein degradation and a lack of protein quality control. Cellular responses to VCP mutant range from aberrant cell cycle regulation, nuclear envelope formation, transcription factor processing, and prevention of polyglutamine aggregation [8, 9]. The VCP protein has four domains: N-terminal domain, C-terminal domain and two ATPase domains, D1 and D2. It has been reported that pathogenic variants in VCP protein induce conformational alterations resulting in significantly perturbed co-factor interactions and altered ATP binding [8, 9]. In vitro assays have shown that VCP pathogenic mutants have enhanced ATPase activity [10–12]. Strategies to suppress the increased ATPase activity of VCP disease mutants or restoring it to baseline are yet to be developed for clinical trials. Mutations in VCP also affect the consolidation of aggregation-prone proteins into inclusion bodies and disrupt the autophagic degradation of ubiquitylated proteins, resulting in the accumulation of non-degradative autophagosomes [13, 14]. Disruption of autophagy due to VCP mutations leads to the buildup of sequestosome-1 (SQSTM/p62) and the activated form of microtubule-associated protein 1A/1B light chain 3B (MAP1LC3B). VCP mutations also lead to the mislocalization and aggregation of TAR DNA-binding protein 43 (TDP-43) [15, 16].

Advancements in elucidating the VCP ATPase structure have significantly enhanced our understanding of its regulatory mechanisms, thus helping to identify drug-like allosteric and ATP-competitive inhibitors bound to the protein [17]. Several VCP inhibitors have been reported, including DBeQ, ML240, NMS-873, CB-5083, and NMS-249. Previous work by our group and others have reported that VCP inhibitors are efficacious in improving VCP-related MSP1 pathology. Zhang et al. (2017) observed that VCP inhibitors potently rescued disease phenotypes in *Drosophila* and patient fibroblasts [18]. Proteomic analysis by Wang et al. (2022) showed that VCP inhibitors reversed RNA processing and cell cycle-related dysregulated proteins.[19]. Harley et al. also raised the possibility of leveraging VCP inhibitors that specifically target the D2 ATPase domain as a therapeutic strategy for VCP-related ALS [20]. Our previous studies using the small molecule allosteric VCP inhibitor CB-5083 in R155H patient iPSC-derived myoblasts, and the R155H knock-in mouse model showed an increase in the expression of autophagic markers and amelioration of muscle weakness, and TDP-43 expression levels [21, 22]. However, we could not proceed with a patient trial in VCP disease because a phase 1 trial of escalating doses of CB5083 was halted in patients with solid tumors and multiple myeloma due to suspected off-target activity which led to visual adverse events [22]. We conducted extensive studies in mice and found that the vision loss is indeed related to inhibition of photoreceptor phosphodiesterase 6 (PDE6), a protein complex family, which is highly concentrated in the retina. The adverse effect of CB-5083 in the retinae in the mice was seen primarily at high doses of 30 mg/kg, and was reversible upon drug discontinuation [22].

Antisense oligonucleotides (ASOs) have become a promising therapeutic modality for genetic disorders, by directly modulating gene expression at the RNA level. [23, 24]. ASOs are short, synthetic nucleic acids, designed to target mRNA by Watson-Crick base pairing, and once bound to the target RNA, can modulate RNA function through multiple mechanisms, including RNase H1-mediated mRNA degradation, and splicing modulation. For diseases related to pathogenic frameshift and nonsense mutations, ASOs have become a top consideration, owing to straightforward validation in disease model systems and demonstrated successes in clinical translation [25, 26]. Leading the way for ASOs currently in medical use is nusinersen which is approved for the treatment of multiple forms of spinal muscular atrophy [27]. Currently approved ASO-based treatments include eplontersen for hereditary transthyretin amyloidosis with polyneuropathy (ATTRv-PN) [28], olezarsen for familial chylomicronemia syndrome (FCS)[29], eteplirsen for Duchene muscular dystrophy (DMD) [30], and tofersen for SOD1-related ALS [31]. It was recently reported that an experimental ASO targeting the FUS transcript (ION363) effectively suppressed both wild-type and mutant FUS expression in the brain and spinal cord of P517L and Δ14 heterozygous mice [32]. Also, targeted ASO-mediated Atp1a2 knockdown in astrocytes reduce SOD1 aggregation and accelerate disease onset in mutant SOD1 mice [33]. Given the Gain-of-function nature of VCP mutations, we hypothesize that VCP reduction ameliorates the clinical manifestations of this debilitating disease by normalizing VCP activity, thus improving the pathology resulting from the disrupted pathways.

In this study, we evaluate if ASO mediated reduction of VCP ameliorates VCP-related proteinopathy phenotypes in human induced pluripotent stem cells-derived skeletal muscle progenitor cells (hiPSC-SMPCs) with the most prevalent VCP R155H mutation, and in the humanized VCP A232E mice [2, 34]. We found that ASO treatment in VCP R155H patient-derived SMPCs and transgenic mouse models of VCP MSP1 significantly reduced VCP expression, improved autophagy flux, and decreased TDP-43 pathogenesis. These results suggest that knockdown of VCP allele in SMPCs and early in asymptomatic mice could be beneficial in preventing progression of the MSP1 myopathy and holds promise for treatment in patients.

## Results

### Successful differentiation of VCP R155H patient-derived iPSC to myogenic lineage

We developed a cellular myopathy model of VCP disease by differentiating hiPSC carrying the VCP p.R155H variants to SMPCs and assessed the efficacy of VCP ASOs in these cells. **Figure 1A** illustrates the timeline and major milestones of hiPSC differentiation and maturation into myotubes. It has been previously reported that directed differentiation of any hiPSC line can be achieved through line-specific tailoring of initial seeding density and addition of the small molecule CHIR99021, an agonist of the Wnt signaling pathway. Indeed, both cell seeding density and Wnt activation play a vital role in mesoderm formation [35, 36]. For VCP hiPSCs, we found that 60,000 cells/cm^2^ and supplementation of 8 µM CHIR99021 for two days sufficiently induced mesoderm and later myogenic differentiation, as noted by the formation of distinct 3D structures one week after mesoderm induction, and later juxtaposed myotube formation (**Figure 1B**). At the completion of directed differentiation, enrichment of skeletal muscle progenitor cells (SMPCs) was achieved by selection of the cell population with the highest expression of the cell surface receptors erb-b2 receptor tyrosine kinase 3 (ERBB3) and nerve growth factor receptor (NGFR) using fluorescence-activated cell sorting (FACS) ([37] (**Figure 1C**). We found that double-positive ERBB3 and NGFR represented only 1.7% of VCP R155H cells, but these enriched SMPCs robustly expressed paired box 7 (PAX7) and sine oculis homeobox homolog 1 (SIX1) **(Figure 1D**). SMPCs could also be induced to form myosin-positive multinucleated myotubes, suggesting successful enrichment (**Figure 1D**) [37]. However, VCP R155H SMPCs displayed reduced myotube fusion efficiency compared to the control SMPCs (P < 0.05) (**Figure 1E**), which may be indicative of their diseased state which we quantified in subsequent figures.

**Figure 1:**
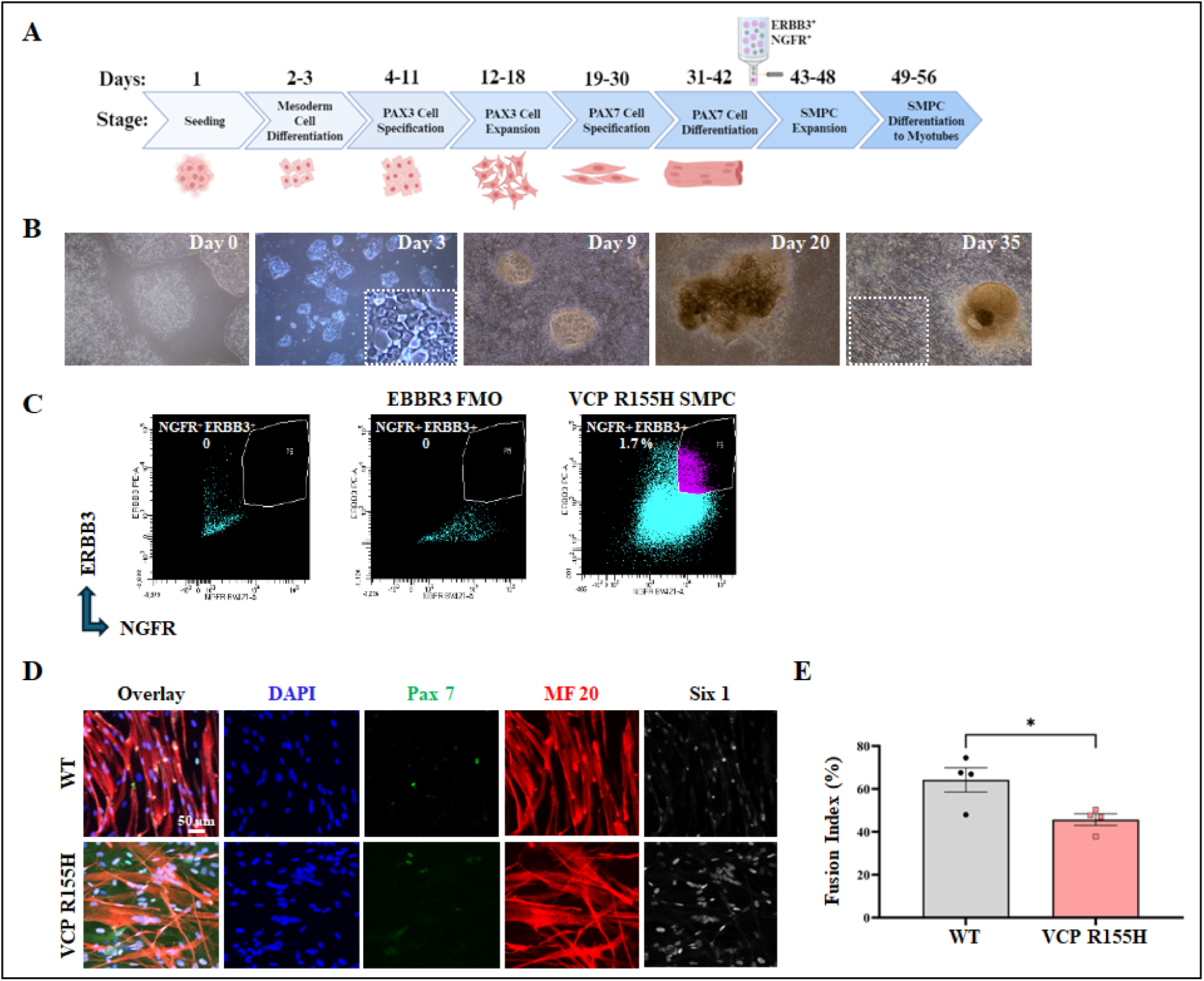
Guided differentiation of patient iPSCs carrying the VCP p.R155H variants to myogenic lineage. (A) Schematic for directed differentiation of hiPSCs to skeletal muscle. Image created with <u>BioRender.com</u>. (B) Representative brightfield images of the key developmental stages during directed differentiation of hiPSCs to myogenic cells show the formation of distinct 3D structures. Inset shows myotube formation. (C) ERBB3+ and NGFR+ skeletal muscle progenitor cells (SMPCs) were sorted at day 42. FACS plots show percentages of SMPCs sorted from VCP lines. (D) Immunofluorescence microscopy of WT and VCP R155H SMPCs differentiated to myotubes show myosin-positive multinucleated myofibers along with SIX1 and PAX7 expression. (E) Fusion index percentage of WT and VCP R155H SMPCs differentiated to myotubes. Statistical analysis was performed using multiple unpaired t test (*p = 0.05).

### Patient-derived SMPCs have TDP-43 mislocalization and elevated levels of autophagic markers

Pathogenic VCP R155H variants are known to lead to the accumulation of ubiquitinated proteins, disruption of autophagy, and impaired retrotranslocation of endoplasmic reticulum-associated degradation (ERAD) substrates [38, 39]. Therefore, we sought to determine the pathophysiological effects of *VCP R155H* pathogenic variants in the patient-derived SMPCs. It has been reported that dysregulation of protein homeostasis due to variants in VCP may cause TDP-43 to undergo cleavage, hyperphosphorylation, and ubiquitination, leading to its accumulation and aggregation in the cytoplasm [15, 40–42]. Phosphorylated TDP-43 (p-TDP-43) is particularly associated with the pathological inclusions seen in TDP-43 proteinopathies. We found TDP-43 and VCP gene expression increased significantly by 1.9 ± 0.5 -fold and 1.9 ± 0.3 -fold, respectively, in VCP R155H cells compared to WT (**Figure 2A**). The expression of the autophagic proteins SQSTM1/p62 and LC3B in VCP R155H cells were 0.3 ± 0.08 -fold and 0.6 ± 0.07 -fold lower, respectively, compared to WT cells, as measured by qRT-PCR analysis **(Figure 2A**). In addition, TDP-43 and p-TDP-43 protein levels were significantly increased by 1.5 ± 0.29 -fold and 2.1 ± 1.32 -fold in VCP R155H cells compared to WT cells (**Figure 2B, C**).

**Figure 2:**
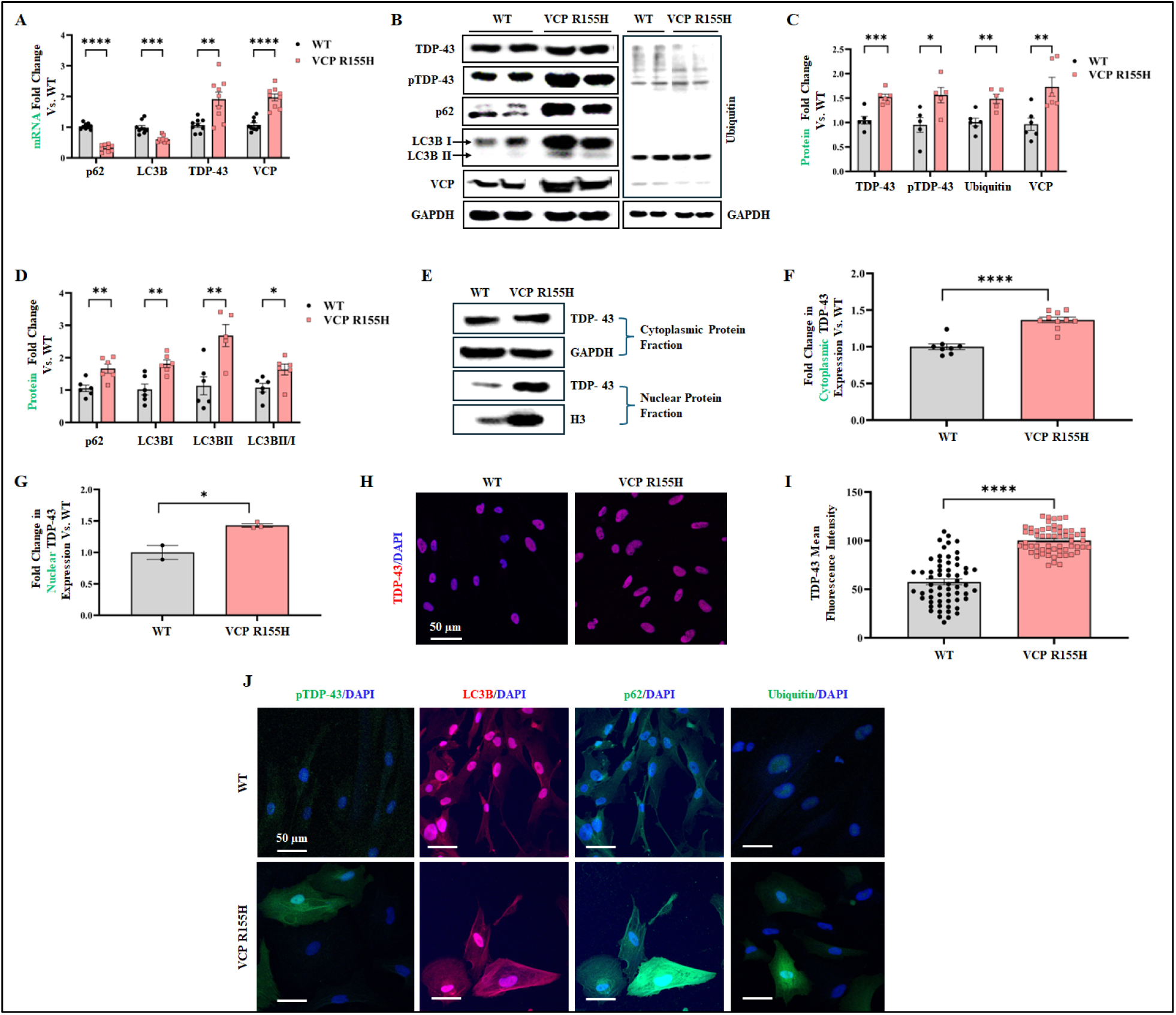
VCP R155H patient iPSC-derived SMPCs show TDP-43 pathology, upregulated VCP expression, and impaired autophagy. (A) Quantitative RT-qPCR analysis of the VCP R155H iPSC-derived SMPCs normalized to wild type (WT) and relative to GAPDH. (B) Western blot analysis of protein expression of SMPCs. GAPDH was used as a positive loading control. (C, D) Densitometry analysis of Western blot of VCP R155H patient iPSC-derived SMPCs normalized to WT and relative to GAPDH. (E) Western blot analysis of cytoplasmic and nuclear fractions of SMPCs. Histone (H3) was used as a positive loading control. (F, G) Densitometry analysis of western blot of (F) cytoplasmic and (G) nuclear fraction of VCP R155H iPSC-derived SMPCs normalized to WT and relative to H3. (H, J) Immunofluorescence microscopy of WT and VCP R155H SMPCs showed increased TDP-43, pTDP-43, SQSTM1/p62, LC3B and ubiquitin positive expression. (I) Mean fluorescence intensity of TDP-43 expression in VCP R155H iPSC-derived SMPCs relative to WT. Statistical analysis was performed using multiple unpaired t tests. *p ≤0.05, **p≤0.01, ***p = 0.001, and ****p = 0.0001 vs WT.

LC3B is a key protein of the autophagy pathway that can be present in two different forms: a cytosolic form (LC3B-I) and a lipidated form (known as LC3B-II), where LC3B-I is conjugated to phosphatidylethanolamine to form LC3B-phosphatidylethanolamine conjugate, which is bound to the autophagosome membrane [43]. The ratio of LC3B-II/I reflect the conversion of LC3B-I to LC3B-II upon autophagy activation. We noted increased expression of LC3B-I by 2 ± 0.73 -fold and LC3B-II by 3.7 ± 2 -fold, respectively in VCP R155H cells compared to WT (**Figure 2B, D**). The LC3 II/I ratio clearly showed enhanced autophagic flux in VCP R155H cells. A 1.7 ± 0.5 -fold elevation in the levels of the autophagic marker SQSTM1/p62 also suggested impaired autophagosome-lysosomal cascade in the VCP R155H cells (**Figure 2B, D**). Increased VCP protein expression by 1.6 ± 0.39 -fold was also observed in VCP R155H cells (**Figure 2B, C**). Through nuclear-cytoplasmic fractionation, we observed significantly elevated cytoplasmic and nuclear TDP-43 levels by 1.37 ± 0.13 -fold and 1.46 ± 0.24 -fold, respectively, in VCP R155H SMPCs, implying TDP-43 mislocalization and accumulation (**Figure 2E-G**). In agreement with the Western blot analysis, immunofluorescence data showed significant increase in nuclear TDP-43 (**Figure 2H, I**), and its cytoplasmic mislocalization through p-TDP-43 (**Figure 2J**). The VCP R155H cells also showed increased SQSTM1/p62, LC3B, and ubiquitin positive expression in the immunofluorescence analysis, therefore consolidating the protein expression data (**Figure 2J**). The VCP R155H SMPCs thus manifest a phenotype characterized by arrested autophagy and TDP-43 mislocalization, providing a useful model to study the effect of ASO treatment in MSP1 disease.

### VCP R155H SMPCs have disrupted lysophagy

VCP plays a crucial role in maintaining lysosomal homeostasis [44, 45]. The disruption of lysophagy due to VCP pathogenic variants leads to lysosomal accumulation, suggesting a defective lysosomal membrane permeabilization (LMP) response in the development of VCP-associated MSP1. VCP R155H and WT cells were subjected to starvation and a combinatorial treatment with bafilomycin A1 (BafA1), an inhibitor of autophagosome-lysosome fusion, to assess autophagic flux and accumulation of autophagosomes [46, 47]. In VCP R155H cells, impaired autophagy led to a significant increase in SQSTM1/p62 protein levels upon BafA1 treatment (**Figure 3A, B**). LC3B-I levels were also significantly higher in BafA1-treated VCP R155H cells compared to WT (**Figure 3A, C**). During autophagic induction, LC3B-II is incorporated into the growing autophagosome membrane, and its level is proportional to the number of autophagosomes in the cell. Western blot analysis revealed that LC3B-II level in WT did not change upon starvation, but was significantly enhanced after BafA1 treatment, indicating activated autophagic flux (**Figure 3A, D**). VCP R155H cells also showed a significant increase in LC3B-II levels upon BafA1 treatment (**Figure 3A, D**). However, treatment with BafA1 resulted in significantly higher level of LC3-II and the ratio of LC3B-II/I in VCP R155H cells compared to WT (**Figure 3E**). After 24 h of recovery, the LC3B-II/I ratio returned to basal levels in WT cells (**Figure 3E**). On the other hand, a higher LC3B-II/I ratio persisted in VCP R155H cells due to disrupted autophagy (**Figure 3E**). In line with the Western blot data, immunofluorescence staining showed that the number of SQSTM1/p62-positive dots was significantly higher in starved VCP R155H cells, and further increased in the presence of BafA1 (**Figure 3F**). Similarly, starvation led to a marked increase in the number of LC3B-positive puncta, which was further amplified with BafA1 treatment in VCP R155H cells (**Figure 3G**).

**Figure 3:**
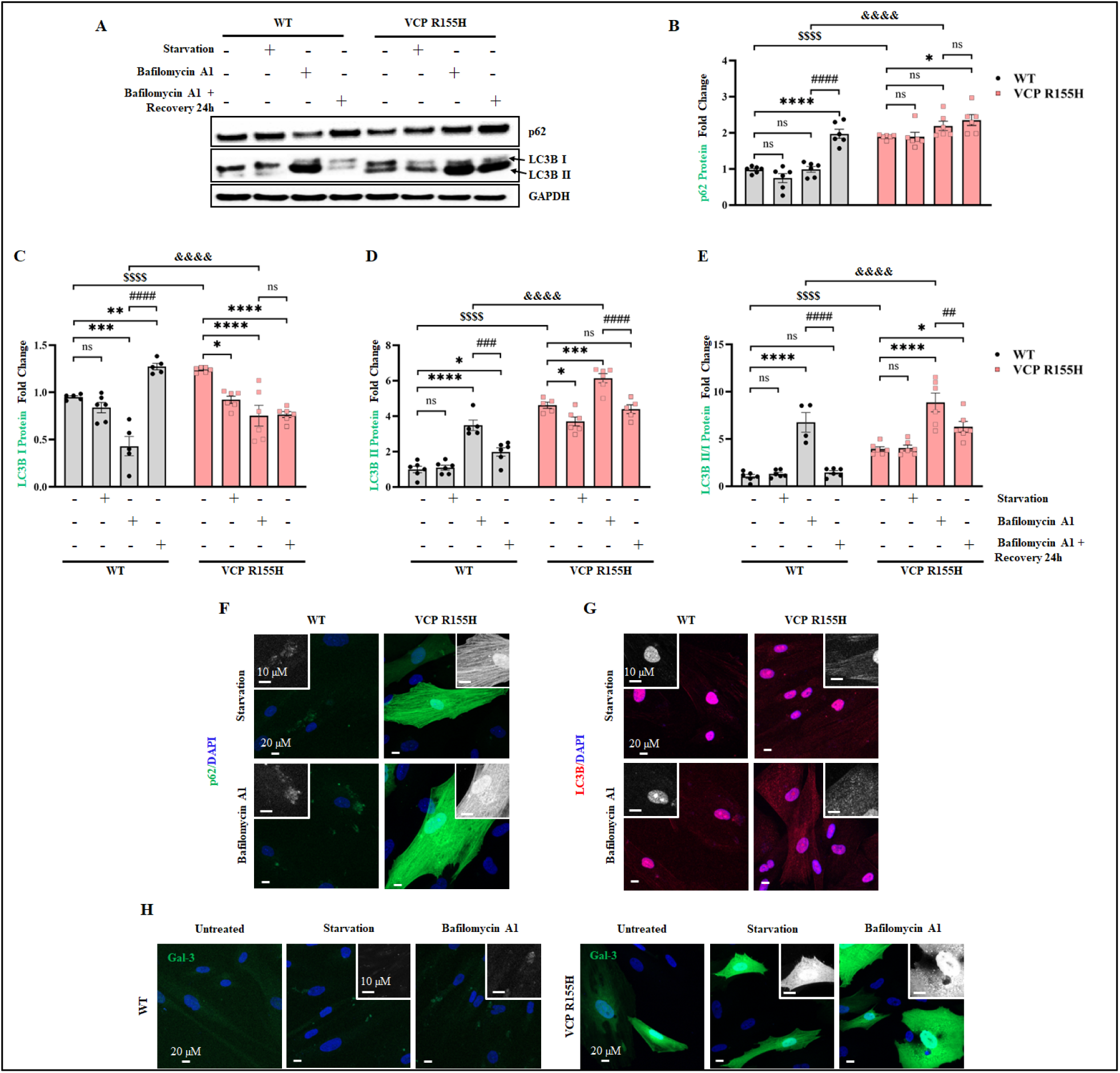
**VCP R155H patient iPSC-derived SMPCs are unable to clear damaged lysosomes**. (A) Western blot analysis of WT and R155H iPSC-derived SMPCs treated with bafilomycin A1 for 3h, followed by a 24h recovery period. (B, C, D, E) Densitometry analysis of Western blot analysis of VCP R155H SMPCs treated with bafilomycin A1 normalized to WT SMPCs and relative to GAPDH. (B) SQSTM1/p62, (C) LC3B-I, (D) LC3B-II, (E) LC3B-II/I. (F, G, H) Immunofluorescence microscopy analysis of WT and VCP R155H SMPCs stained with (F) SQSTM1/p62, (G) LC3B, (H) Galectin-3. Inset represents magnified image of the selected areas. Statistical analysis was performed using two-way ANOVA followed by Šídák’s multiple comparisons test. *p ≤0.05, **p≤0.01, ***p = 0.001, and ****p = 0.0001 vs untreated WT/ VCP R155H); ## p≤0.01, ### p = 0.001 and #### p = 0.0001 vs starvation (WT and VCP R155H); $$$$ p = 0.0001 WT vs VCP R155H untreated; &&&& p = 0.0001 bafilomycin A1 treatment corresponding WT vs VCP R155H

Lysosomal damage in VCP R155H cells was further examined by analyzing galectin-3 (Gal-3), a β-galactoside-binding lectin that is recruited to the lysosomal membrane solely upon membrane permeabilization. In WT cells under basal conditions, Gal-3 exhibited a diffuse cytoplasmic distribution with no detectable puncta (**Figure 3H**); however, starvation-induced lysosomal damage resulted in the translocation of Gal-3, and its consequent binding to compromised lysosomes, forming distinct puncta on the damaged lysosomal membrane. We observed that VCP R155H cells expressed higher Gal-3 puncta in the basal state. Upon starvation, the number of Gal-3 puncta increased in both WT and R155H cells due to lysosomal damage. However, compared to WT, the number of Gal-3 puncta in VCP R155H cells was significantly higher, and it further increased upon BafA1 treatment. These results show that VCP R155H patient cells have alterations in lysosomal stability and dynamics.

### VCP ASO modulated TDP-43 pathology and autophagic markers in VCP R155H SMPCs

To investigate the therapeutic potential of VCP ASO in rescuing disease phenotypes in VCP R155H cells, we first examined the cytotoxicity of ASOs on the VCP R155H SMPCs at doses ranging from 50 nM to 5 µM. A cell survival rate of ≥ 98 % was measured in cytotoxicity studies where VCP R155H SMPCs had been treated with ASO up to 5 µM (**Figure 4A**). No significant differences were observed between control ASO and VCP ASO treatment. The VCP ASO was designed to specifically knock down VCP mRNA using RNase H1-mediated degradation. Indeed, treatment with 1.2 µM VCP ASO reduced VCP protein expression by 50% (95% CI [19–68]) in the cells, whereas control ASO demonstrated no change in VCP protein levels (**Figure 4B**). Based on this promising result, we sought to further explore the relationship between different levels of VCP reduction and improvements in disease traits by conducting a dose-response experiment. The SMPCs were treated with four different concentrations of ASOs: 0.3 µM, 0.6 µM, 0.9 µM, and 1.2 µM, to correlate the level of ASO-mediated VCP mRNA and protein reduction with various molecular signatures that are characteristic of MSP1 disease pathology (**Figure 4C-K**). qRT-PCR analysis and Western blot analysis revealed that ASO treatment resulted in 85% (95% CI [58–93]) reduction in VCP mRNA level and 48% (95% CI [39–56]) reduction in VCP protein expression, respectively, with no significant changes between the different ASO concentrations (**Figure 4C, E, F**). We observed increased LC3B mRNA and ratio of LC3B-II/I protein expression upon ASO treatment, which may suggest activation of autophagic flux (**Figure 4D, K**). Furthermore, VCP ASO treatment resulted in a declining trend (not significant, except for 1.2 µM) of SQSTM1/p62 protein (**Figure 4H**), despite an increase in its mRNA levels (**Figure 4D**), which is in agreement with the notion that SQSTM1/p62 is degraded via autophagy. TDP-43 protein expression decreased 60% (95% CI [34–58]) upon ASO treatment compared to untreated cells (**Figure 4G**). Immunofluorescence confirmed the ASO mediated knockdown of VCP expression in VCP R155H cells and associated reduction in pTDP-43 levels **(Figure 4L)**. Consistent with the Western blot data, the immunofluorescence analysis also showed that VCP ASO treatment resulted in upregulation of LC3 and concomitant downregulation of SQSTM1/p62 protein levels, indicating initiation of autophagy (**Figure 4M**).

**Figure 4:**
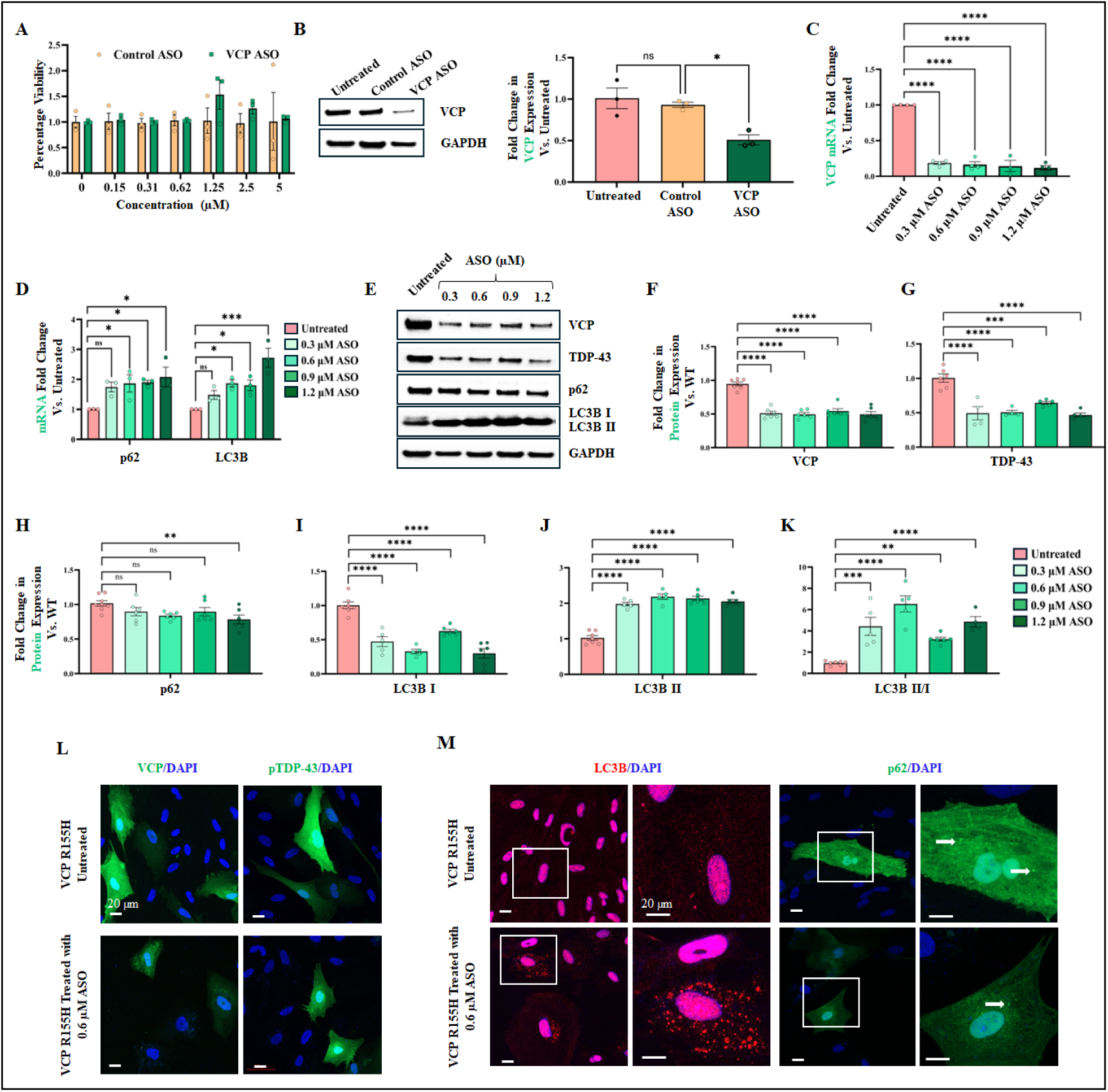
hVCP knockdown by ASO and associated effects in VCP R155H SMPCs. **(A)** Cell cytotoxicity assay by MTT showed good cell viability up to 5 µM VCP ASO. **(B)** Western blot and corresponding densitometric analysis to quantify VCP protein levels in iPSC-derived SMPCs treated with 1.2 µM control ASO or VCP ASO. **(C-K)** Dose-responsive effects of ASO on various molecular hallmarks of MSP1 pathology in patient VCP R155H iPSC-derived SMPCs. **(C)** qRT-PCR for VCP**, (D)** qRT-PCR for SQSTM1/p62 and LC3B, **(E)** Western blot analysis for VCP, TDP-43, SQSTM1/p62, LC3B-I, LC3B-II, **(F-K)** Densitometry analysis of Western blots in E reports the protein levels of **(F)** VCP, **(G)** TDP-43, **(H)** SQSTM1/p62, **(I)** LC3B-I, **(J)** LC3B-II, **(K)** LC3B-II/I relative to untreated VCP R155H cells, following normalization by GAPDH. **(L, M)** Representative confocal images of immunofluorescence staining in VCP R155H SMPCs, either untreated or treated with 0.6 µM VCP ASO. White arrows indicates the presence of SQSTM1/p62 puncta in SMPCs. Statistical analysis was performed using one-way ANOVA followed by Dunnett’s multiple comparisons test. *p ≤0.05, **p≤0.01, ***p = 0.001, and ****p = 0.0001 vs untreated VCP R155H.

### VCP ASO restores lysophagy function in VCP R155H SMPCs

Studies have shown that VCP inhibition in mutant VCP cells rescues defective lysophagy [48]. We utilized VCP ASO to accelerate clearance and prevent the persistence of damaged lysosomes in VCP R155H cells. We observed a decrease in SQSTM1/p62 protein expression upon ASO treatment (**Figure 4H**). To investigate whether p62 degradation was mediated by autophagy, VCP R155H cells were co-treated with 0.6 µM VCP ASO and Baf A1 (**Figure 5A, B**). This combination inhibited p62 clearance, which was initially observed with 0.6 µM VCP ASO alone (**Figure 4H**). Furthermore, p62 expression returned to baseline levels 24 hours after Baf A1 washout in VCP R155H cells treated with VCP ASO, implicating autophagic contribution to p62 degradation. Additionally, we noted that the increased expression of LC3B-II following 0.6 µM VCP ASO treatment was further elevated when autophagosome degradation was blocked by Baf A1 treatment (**Figure 5A, C**). Immunofluorescence staining also showed BafA1 treatment increased the number of SQSTM/p62 dots in VCP R155H cells treated with 0.6 µM VCP ASO, suggesting blockage in p62 degradation (**Figure 5D**). At basal level, in VCP R155H cells treated with 0.6 µM VCP ASO, Gal-3 showed uniform cytoplasmic localization without detectable puncta, signifying the possible removal of damaged lysosomes through autophagy (**Figure 5E**). VCP R155H cells without ASO treatment had higher Gal-3 puncta (**Figure 5E**). Upon starvation and BafA1 treatment, the number of Gal-3 puncta increased in both treated and untreated VCP R155H cells (**Figure 5E**). VCP being a key regulator of lysophagy, its targeted inhibition by VCP ASO in VCP R155H cells can potentially rescue lysophagic defects.

**Figure 5:**
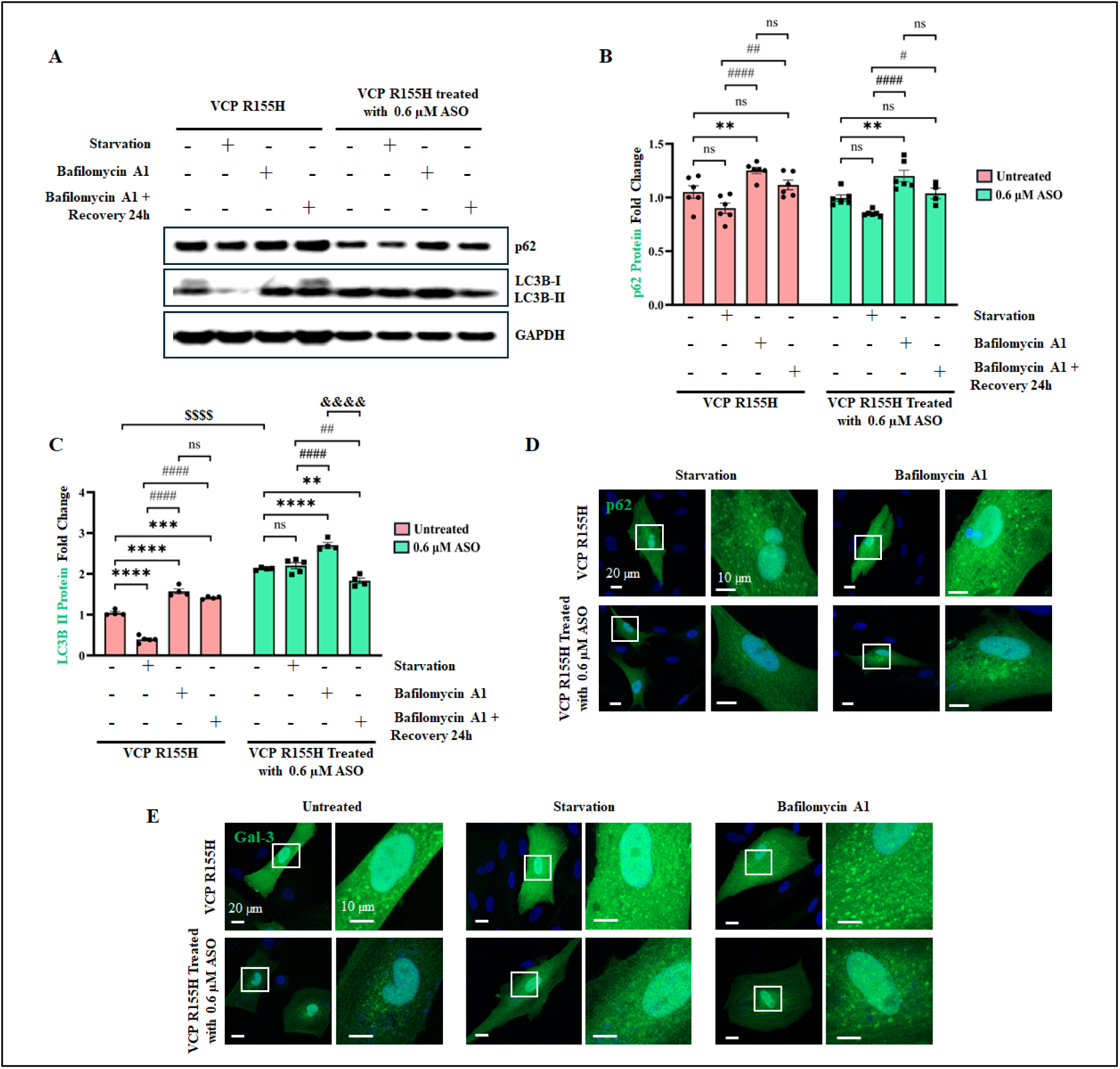
VCP ASO treatment restores lysophagy function in VCP R155H SMPCs. **(A)** Western blot analysis of R155H iPSC-derived SMPCs with and without treatment with 0.6 µM VCP ASO for 7-days, followed by 3h bafilomycin A1 treatment and a 24h recovery period. (**B**, **C**) Densitometry analysis of Western blot analysis of VCP R155H SMPCs treated with 0.6 µM VCP ASO followed by bafilomycin A1 treatment normalized to untreated VCP R155H SMPCs and relative to GAPDH. (**B**) SQSTM1/p62, (**C**) LC3B-II, (**D, E**) Immunofluorescence microscopy analysis of VCP R155H SMPCs with and without treatment with 0.6 µM VCP ASO followed by bafilomycin A1 treatment, stained with (**D**) SQSTM1/p62, (**E**) Galectin-3. Statistical analysis was performed using two-way ANOVA followed by Šídák’s multiple comparisons test. *p ≤0.05, **p≤0.01, ***p = 0.001, ****p = 0.0001 vs VCP R155H SMPCs (with and without VCP ASO); # p ≤0.05, ### p = 0.001 and #### p = 0.0001 vs starvation in VCP R155H SMPCs (with and without VCP ASO); $$$$ p = 0.0001 VCP R155H with-vs without VCP ASO; &&&& p = 0.0001 bafilomycin A1 treatment in VCP R155H with vs without VCP ASO.

### VCP ASO improves motor function in A232E mice

We conducted *in vivo* studies using a transgenic humanized mouse model of VCP disease harboring the most severe mutation, VCP A232E (gifted by Paul Taylor) [49]. To initially generate transgenic mice expressing normal human VCP in all tissues, human VCP cDNA was placed under the control of the CMV-enhanced chicken beta-actin promoter permitting widespread VCP expression in muscle, brain, and bone. The disease-associated mutation A232E was then introduced by site-directed mutagenesis. The resulting transgenic animals were backcrossed onto the C57BL/6 background. The translational value of the VCP A232E transgenic mouse model has been characterized in previous studies [49]. The mice exhibited progressive muscle weakness; histological analyses of the quadricep muscle revealed signs of myogenic myopathy, such as irregular fiber size, centrally located nuclei, and inflammatory cell infiltration; and histopathological analysis of the brain demonstrated ubiquitin-positive cytoplasmic accumulations of TDP-43. We additionally performed a natural history study in these mice in our laboratory and found differences between WT and VCP A232E mice in functional studies, including Rotarod, grip strength, and inverted screen. Biochemical analysis of quadriceps, tibialis anterior, and diaphragm muscle tissue revealed TDP-43 pathogenesis and impaired autophagy similar to the previous report [49].

To ensure the absence of ASO-related toxicity in the mouse studies, 10-month-old VCP A232E mice were dosed with either control or VCP ASOs at 50 mg/kg, once a week for 12 weeks, via subcutaneous injection. Age and sex-matched WT mice were used as a reference control (**Figure S1A**). At the end of the 12-week dosing regimen, liver, spleen, and kidneys were collected from the mice, and their weights were measured and analyzed. A significant increase in organ weights normalized to body weights for VCP ASO-treated mice was considered a sign of potential toxicity. The serum levels of the liver transaminases aspartate aminotransferase (AST) and alanine aminotransferase (ALT), as well as the levels of creatine kinase (CK), whose elevation (more than 2-fold) would indicate concerning liver or muscle toxicity, were also measured (**Figure S1B-H**). We observed no significant changes in organ weights or serum parameters between VCP ASO and control ASO treatment groups at 12 weeks following injections (**Figure S1B-H**). Therefore, we conclude that under the experimental conditions used, the VCP ASO was well tolerated in mice. Motor strength and coordination were assessed monthly in VCP A232E mice that received subcutaneous injections of 50 mg/kg VCP ASO or control ASO once a week from 6 to 9 months of age (**Figure 6A**). A group of age-matched untreated WT mice was included in the study as reference controls. Motor performance in the different groups of mice was evaluated with the Rotarod, grip strength, and inverted screen assays, and both raw data and percent change from the respective baseline values were analyzed. VCP A232E mutant mice dosed with VCP ASO showed significant improvement in Rotarod performance compared to control ASO-treated A232E mice at 2 months of treatment (**Figure 6B**). Motor testing data trended for improved performance in the inverted screen test with VCP ASO treatment in A232E mice, but did not reach statistical significance (**Figure 6D**). Percent change analysis of rotarod and inverted screen tests showed significant improvement in VCP ASO treated mice after 2 and 3 months of treatment when compared to the control ASO treated mice **(Figure 6C, E).** There was no change in grip strength performance (**Figure 6F, G**).

**Figure 6.**
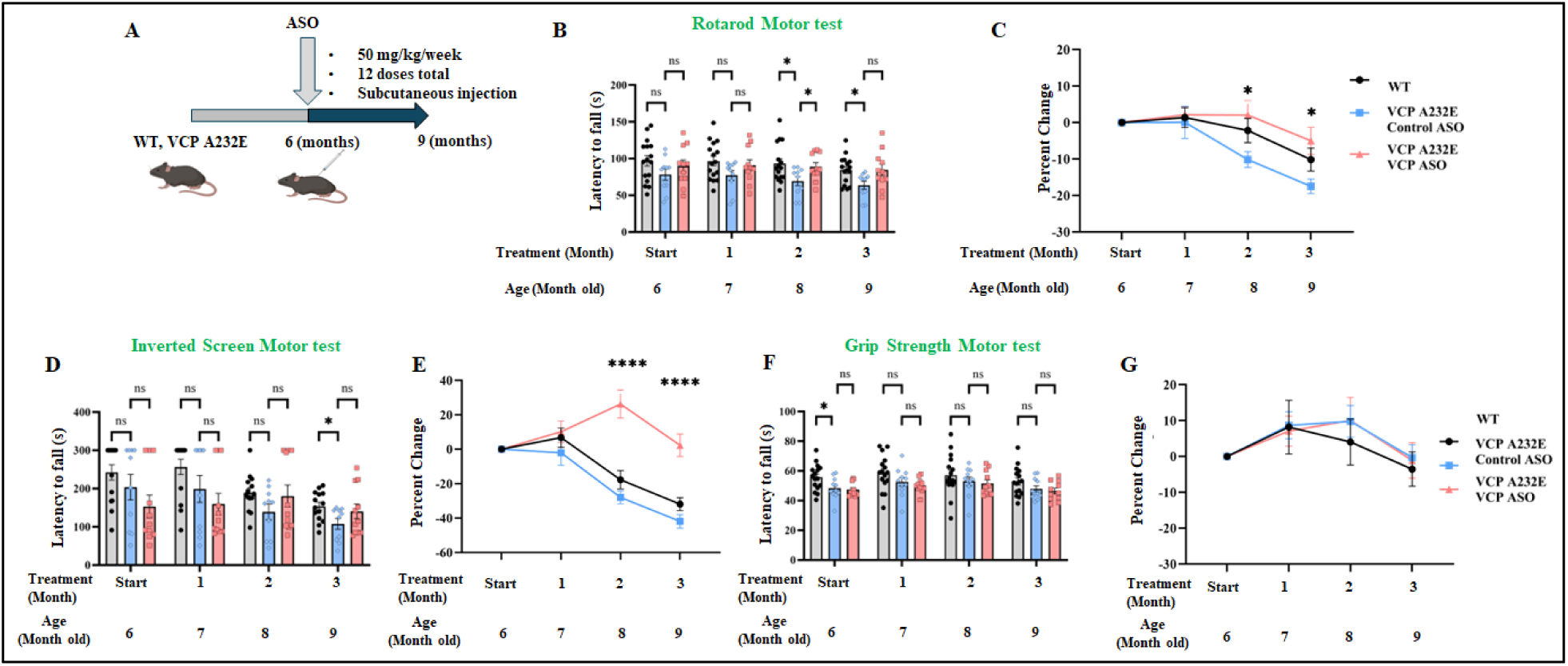
Motor testing analysis of WT and VCP A232E mice treated with VCP ASO and control ASO. **(A)** Schematic illustration of experimental design. 6-month-old VCP A232E mice (n=11-16) were dosed weekly by subcutaneous injection with control ASO, or VCP ASO (50 mg/kg/week) for a total of 3 months. **(B, D, F)** Rotarod, inverted screen testing, and grip strength, in WT mice and A232E mice treated with either control ASO or VCP ASO. Testing started at 6 months of age and was performed monthly for 3 months. **(C, E, G)** Raw data was analyzed in addition to percent change from baseline testing in the different groups and normalized with control ASO group for comparison. **(C)** Percent change in Rotarod, **(E)** Percent change in inverted screen, and **(F)** Percent change in grip strength. Statistical analysis was performed using two-way ANOVA followed by Dunnett’s multiple comparisons test. *p ≤0.05, and ****p≤0.0001 vs control ASO ns: not significant.

### VCP ASO improves myopathy and biochemical abnormalities in the A232E mouse model of MSP1 disease

Taylor et al. (2010) reported that VCP A232E mice muscle exhibit pathological signs of myogenic myopathy, such as fiber size irregularity, centralized nuclei, inflammation, and TDP-43-positive intranuclear and cytoplasmic aggregates [49]. Consistent with those reports, we observed that 9-month-old VCP A232E mice treated with control ASO had significantly higher TDP-43 mRNA (2.94 ±1.4 -fold elevation) and protein (1.3 ± 0.07 -fold increase) expression in quadriceps muscle compared to WT littermates. Treatment with VCP ASO significantly reduced TDP-43 mRNA levels by 0.23 ± 0.13 -fold and protein expression by 0.75 ± 0.03 -fold compared to control ASO in quadriceps muscle of VCP A232E mice (**Figure 7A, B, D)**. SQSTM1/p62 and LC3B mRNA expression were 4 ± 0.98 -fold and 6.3 ± 1.6 -fold higher, respectively in quadriceps muscle of VCP A232E mice treated with control ASO compared to WT (**Figure 7A**). Treatment with VCP ASO decreased SQSTM1/p62 and LC3B mRNA expression by 0.3 ± 0.08 -fold and 0.09 ± 0.04-fold, respectively compared to control ASO treatment **(Figure 7A)**. Similarly, SQSTM1/p62 and LC3B protein levels were reduced 22% (95% CI [13–31]) and 45% (95% CI [32–55]) following VCP ASO dosing in quadriceps muscle of VCP A232E mice compared to control ASO **(Figure 7B, E-G)**. Additionally, VCP ASO dosed for 12 weeks at 50 mg/kg via subcutaneous injection resulted in 0.45± 0.09-fold knockdown of VCP mRNA and a 30% (95%CI [27–32]) reduction in VCP protein expression compared to control ASO in quadriceps muscle of VCP A232E mice **(Figure 7A-C)**. Hematoxylin and eosin (H&E) staining of histological sections of quadriceps muscle from VCP A232E mice dosed with VCP ASO showed improved myofiber morphology compared to control ASO (**Figure 7H**). Indeed, WT mice had homogeneously sized myofibers with peripherally located nuclei, whereas 9-month-old VCP A232E mice dosed with control ASO showed cellular infiltrations into the interstitium of the myofibers along with a significantly increased number of centralized nuclei, consistent with MSP1 myopathy. VCP ASO treatment significantly decreased the number of central nuclei compared to control ASO (control ASO: 34 ± 2% myofibers had centralized nuclei, VCP ASO: 20 ± 2% myofibers had centralized nuclei; over 150 myofibers examined from 3 mice per group) (**Figure 7I**).

**Figure 7:**
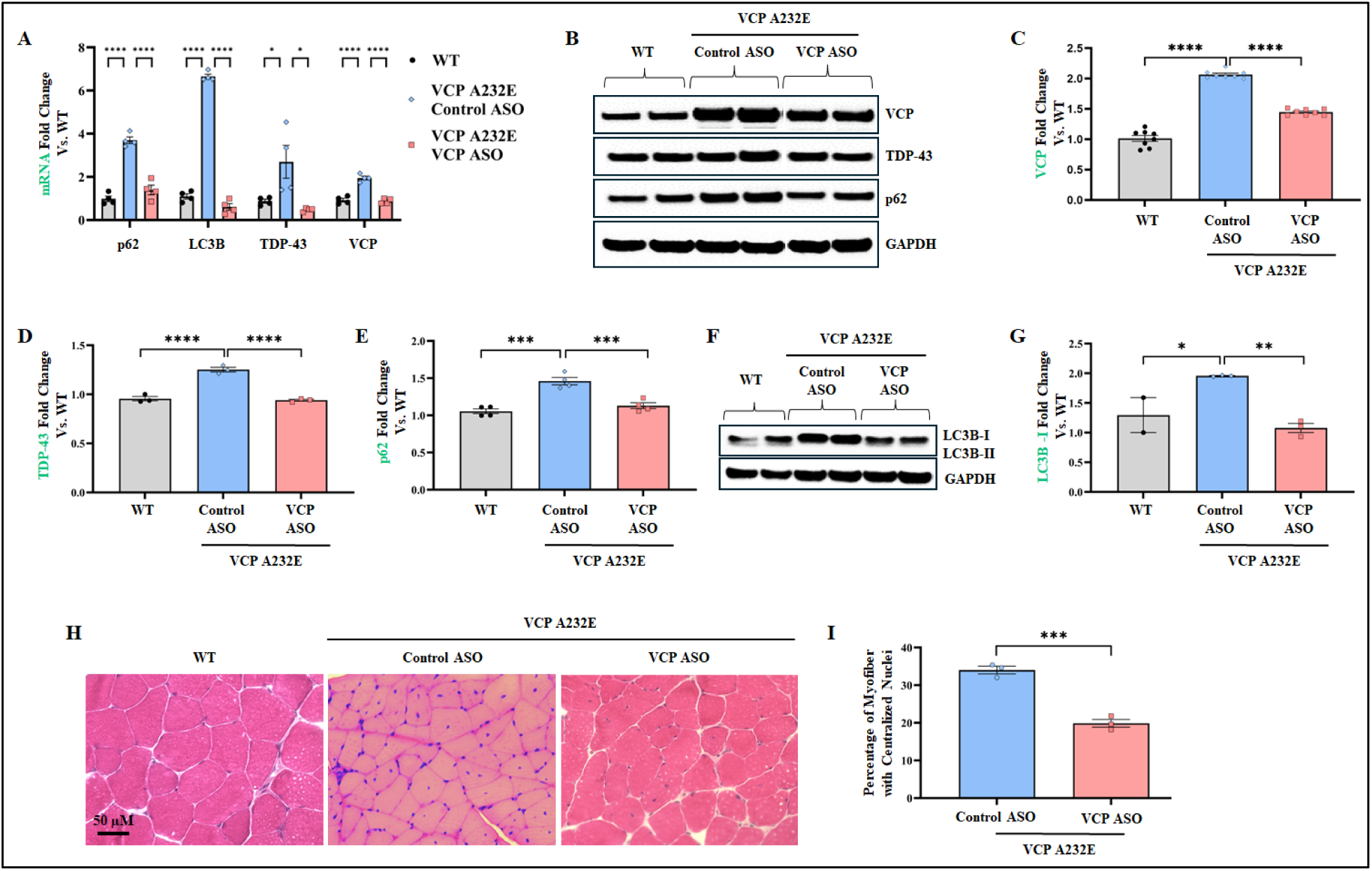
Improvement of disease hallmarks in VCP A232E mice dosed with VCP ASO. **(A)** VCP ASO treatment reduced VCP mRNA levels in quadriceps. **(B, F)** Western blot analysis of VCP and other proteins in the autophagy pathway. **(C, D, E, G)** Densitometric analysis of Western blot shown in B. **(C)** VCP, **(D)** TDP-43, **(E)** p62, **(G)** LC3B-I. **(H)** H&E staining of myofibril structure in histological sections of quadriceps muscle collected from WT or A232E mice dosed with control ASO or VCP ASO at the end of the 3-month study. **(I)** Quantification of the centralized nuclei in the histological sections of quadricep muscle from VCP A232E mice dosed with 50 mg/kg ASO for 3 months. One-way ANOVA test followed by Dunnett’s multiple comparisons test was used for qPCR and WB densitometry quantification. Unpaired t test was used for central nuclei quantification (n=3 mice per group). *p = 0.05, **p = 0.01, ***p = 0.001, and ****p = 0.0001 vs control ASO. (n = 3 mice per group).

## Discussion

VCP autosomal dominant multisystem proteinopathy caused by gain-of-function missense variants is associated with different levels of hyperactive enzymatic activity, which in turn are associated with clinical variability in the phenotype [10–12]. To ameliorate the gain-of-toxicity, an ideal approach is to reduce the expression of VCP mutant proteins. While previous reports have indicated that small molecules improve the phenotype in preclinical studies in drosophila, patient derived myoblast cells, and in the VCP R155H mouse model [18, 21, 22], the compounds used were not approved for patient trials requiring a long duration of treatment for this chronic disease. A phase I trial for cancer was terminated because of adverse effects on vision, such as photophobia and dyschromatopsia [22]. To address this shortcoming, we identified a potent ASO targeting VCP RNA that safely knocks down human VCP expression by virtue of Rnase H1-mediated mRNA degradation. ASO treatment in functionally validated iPSC-derived VCP R155H SMPCs and transgenic mouse models of VCP disease significantly reduced VCP expression, improved autophagy markers, and decreased TDP-43 pathogenesis. Our study provides proof of principle that reducing the expression of VCP using ASOs dilutes the toxic effect of the protein, restores homeostasis, and ameliorates VCP-associated disease pathology. Based on our findings we propose that VCP-targeting ASOs could be a therapeutic option in downregulating VCP in patients.

ASOs are uniquely tailored to specifically correct the root cause of the disease at the RNA level, offering a strategic alternative therapeutic strategy over treatments focused on downstream processes[24]. Many antisense drug candidates are presently in clinical trials for a broad spectrum of diseases, including endocrine, metabolic, cardiovascular, neurological, inflammatory, neuromuscular, and infectious disorders[24]. Currently, no antisense therapeutics have been approved by the FDA for the treatment of VCP disease-associated MSP1. Additionally, as the VCP function is indispensable for cellular functions, therapeutic strategies for MSP1 should ideally focus on selective modulation of VCP activity, balancing efficacy with safety.

We assessed the effects of ASOs, specifically targeting the human VCP gene in the patient iPSC-derived SMPCs bearing the most common VCP R155H/+ disease variant and the humanized A232E/+ mouse model. The VCP R155H and A232E mutation, located in the N domain and D1 ATPase domain, respectively, destabilize the interaction between the N and D1 domains through steric hindrance, interfering with VCP’s ability to interact with substrates and cofactors, thereby compromising protein quality control mechanisms[10]. In our laboratory, patient iPSC-derived SMPCs were characterized and were shown to have higher protein expression levels of the autophagy markers LC3-II/I, SQSTM1/p62, VCP, and TDP-43, as well as translocation of TDP-43 to the cytoplasm, as previously reported [21, 50]. These phenotypic alternations make them a suitable model to assess the therapeutic potential of ASOs.

The response to lysosomal damage is emerging as a critical cellular mechanism for maintaining homeostasis, and its dysfunction may play a role in VCP disease pathogenesis [51, 52]. Efficient lysosome turnover relies on a coordinated sequence of events, including the VCP-dependent removal of ubiquitylated substrates [53]. The absence of functional VCP may disrupt the clearance of ubiquitinated substrates from compromised lysosomes and initiation of autophagy. We found that bafilomycin A treatment, which disrupts the lysosomal degradation pathway, causes accumulation of SQSTM1/p62 and increased LC3B-I to LC3B-II conversion in R155H VCP cells compared to WT. We further observed that VCP mutation disrupts the clearance of Gal-3-positive damaged lysosomes in VCP R155H cells. Other studies have also reported R155H VCP mutations lead to acute lysosomal membrane permeabilization and impair the cell’s ability to repair damaged lysosomes [48, 54].

We showed that ASO treatment was effective in targeting VCP mRNAs and reduced VCP protein expression by 0.6-fold in SMPCs compared to untreated SMPCs. Zhang et al. reported that the R155H mutation causes ∼ 2-fold increase in VCP’s ATPase activity[55]. A decrease in VCP expression upon ASO treatment would consequentially reduce VCP’s ATPase activity; however further studies are needed to determine the fold decrease in VCP’s ATPase activity. In human motor neurons, enhanced D2 ATPase activity of mutant VCP was reported as the contributing factor to TDP-43 mislocalization [20]. ML240 and CB-5083 selective VCP inhibitor reversed the mislocalization of TDP-43 by pharmacologically inhibiting the D2 ATPase domain of VCP protein [20]. In agreement, we found VCP ASO decreased TDP-43 nuclear-to-cytoplasmic mislocalization and aggregation seen in VCP R155H cells. Corroborating earlier findings, we observed a steady upregulation of autophagy-related markers following VCP knockdown [56]. Lee et al. also suggested that silencing VCP via siRNA increases the LC3B-II/I ratio, and this increase was further enhanced following BafA1-mediated inhibition of autophagosome degradation [57].

We observed no significant adverse effects from VCP ASO administration in mice. Additionally, organ analysis confirmed no signs of local or peripheral toxicity in the liver, spleen, and kidney. Muscle pathology assessments showed that the characteristic abnormalities in VCP A232E mice were substantially reduced following VCP ASO treatment. Targeting of VCP mRNA by the ASO resulted in widespread downregulation of VCP protein expression in the quadriceps muscle of VCP A232E mice.

ASOs have emerged as a promising RNA-targeting strategy for neurological diseases, represented by the recent FDA approval of Tofersen, an ASO designed to reduce SOD1 mRNA in ALS patients carrying SOD1 mutations. Tofersen demonstrated improvements in disease biomarkers by lowering production of the toxic SOD1 protein [58]. In our results, ASO treatment led to a reduction in TDP-43 pathology, which was accompanied by improved behavioral outcomes in VCP A232E mice, highlighting the therapeutic potential of targeting TDP-43 dysregulation in multisystem proteinopathy. This is consistent with prior studies in ALS/FTD models showing that ASO-mediated suppression of TDP-43 improves behavioral dysfunction, indicating their therapeutic potential for TDP-43-related disorders [59]. In contrast to our in vitro data, in vivo treatment of VCP ASO led to reduction in autophagy markers SQSTM1/p62 and LC3B. We hypothesize that the 3-month ASO regimen mitigated protein aggregation, leading to a decreased necessity for autophagy activation in vivo. The molecular changes induced by the ASO treatment thus led to improvement of motor function in A232E mice. Studies have shown that blocking ATP hydrolysis not only counteracts the downstream effects of excessive hydrolysis but also intensifies dominant-negative impacts on ATPase function [20]. Considering the wide functional scope of VCP, it is plausible that VCP mutations not only exert dominant gain-effects but may also exert loss of function effects, influenced by specific cofactor associations and downstream signaling cascades.

In summary, our study indicates that VCP-targeting ASOs are a safe and potentially efficacious treatment for mitigating muscle pathology in VCP-associated disease. Considering the dose-dependent disruption of cellular homeostasis by VCP inhibitors, future research should focus on developing and optimizing allele-specific ASOs that selectively target the mutant allele while preserving the normal allele. The advancement of VCP inhibitors into phase I clinical trials for cancer, acknowledge their potential for other debilitating and presently incurable disorders.

## Materials and Methods

### Differentiation procedure of iPSCs to SMPCs

The patient iPSCs carrying the VCP p.R155H variants were differentiated to SMPCs using a previously published protocol [35]. Briefly, iPSCs were cultured on mTESR^+^ for at least three passages. After pre-treating the iPSCs for 1h with 10 μM rock inhibitor (RI, Stem cell Technologies) supplemented media, cells were dissociated, seeded at a concentration of 550,000 cells/well (based on our standardization protocol) on Matrigel-coated plates, and grown in mTeSR^+^ with RI media. On days 2 and 3, cells were treated with 8 μM CHIR (Tocris Biosciences) supplemented in E6 medium. Afterwards, the culture medium was changed, and cells were cultured in E6 medium until day 11. Cells were then switched to StemPro-34^+^basic fibroblast growth factor media for an additional 7 days. The culture medium was then changed to E6 medium supplemented with IGF1 for 7 days. The cells were then fed DMEM/F12 with N-2 Supplement (Gibco), Insulin-Transferrin-Selenium (ITS, Gibco), and IGF1 media for 5-7 days, and thereafter SB431542 (Sigma) was added for an additional 7 days. Fluorescence activated cell sorting (FACS) was then conducted using mouse anti-human fluorochrome-conjugated monoclonal antibodies of erbB3/HER-3 (Biolegend, 324705) and NGFR (CD 271, BD Pharmingen™ 562123). Briefly, the cells were dissociated as single cell, incubated in human Fc block antibody to block non-specific binding of Fc, and subsequently fluorochrome-conjugated antibodies were added according to previously published protocol [35]. After 45 min incubation, the cell–antibody conjugates were washed with FACS buffer (2% FBS in sterile PBS) and analyzed with BD ARIA II (BD Biosciences) flow cytometer and analyzed with BD FACSDiva 8.0.2 software. Matched fluorophare-conjugated isotype control antibodies were used to set control gates. The sorted SMPCs were expanded in SMPC expansion media SkGM-2 Bullet Kit (Lonza) for characterization and further experimentation. The SMPCs were differentiated to myotubes by seeding them on Matrigel-coated plates and culturing them in DMEM/F12 medium supplemented with N-2 Supplement, ITS, IGF1, and SB431542 for 5-7 days. To evaluate the differentiation and myotube formation ability of SMPC, cultures were immunostained with MF 20 and PAX7 antibodies. The number of nuclei contained within each myotube was counted and the fusion index was calculated as the percentage of nuclei in myotubes over total nuclei[37]. At least three random images were taken per well, and nuclei were counted using ImageJ.

### Identification of antisense oligonucleotides (ASOs) targeting human VCP

The ASOs used in the studies described here were 16 nucleotides in length with a phosphorothioate (PS) backbone and three 2′-constrained ethyl (cEt)-modified nucleotides at both ends (3-10-3 gapmer configuration). The ASOs were either unconjugated or conjugated to a lipid (C16, palmitate) at the 5’ end via a phosphodiester linkage. A well-characterized control ASO which does not hybridize to any mouse mRNA sequence was included in the experiments. The oligonucleotides were synthesized and purified as previously described [60]. Of the 150 ASOs targeting VCP that were screened and selected, 10 ASOs were assessed *in-vivo* for tolerability by subcutaneous administration in 7-week-old wild-type CD-1 male mice at a dose of 50 mg/kg weekly for 4 weeks and then sacrificed 72 hours after the last dose (data not shown). The best performing ASO was chosen for further experiments.

### MTT assay

Cell cytotoxicity upon ASO treatment was measured using the 3-(4,5-dimethylthiazol-2-yl)-2,5-diphenyl-2H-tetrazolium bromide (MTT) assay (Sigma). Briefly, 1×10^4^ SMPCs per well were seeded in a 96-well plate and treated with control ASO or VCP ASO at concentrations ranging from 50 nM to 5 µM for 48 hours. The treatment was conducted in serum-reduced optiMEM medium overnight. The culture medium was then replaced with complete SMPC medium the following day. Cells were then incubated with 5mg/mL MTT for 3 h at 37 °C. 100 μl of DMSO were added to each well and the culture plate placed on a shaker for 30 minutes to dissolve the formazan crystals, resulting in the formation of a colored solution. The amount of formazan produced in each well was quantified by measuring the absorbance of the colored solution at a wavelength of 570 nm. Three independent replicate measurements were collected and reported as the mean value ± standard deviation.

### ASO treatment of SMPCs

SMPCs were transfected with 300, 600, 900, and 1200 nM ASOs using lipofectamine 3000 (Thermofisher Scientific). Transfection was done in serum-reduced OptiMEM medium overnight and replenished with complete SMPC medium the following day. Cells were treated with ASOs for 7 days where transfection was performed on day 1, day 3, and day 5. After 7 days of treatment, the cells were either fixed for immunocytochemistry or lysed for total RNA and protein extraction. To test autophagy flux, cells were incubated in serum-free medium for 24 hours to induce starvation, then treated with 10 nM bafilomycin A1 (Sigma-Aldrich, B1797) for 3 hours prior to harvesting, following previously published protocol [61]

### RT-qPCR

Total RNA was extracted from SMPCs, ASO-treated SMPCs, and mouse tissues using the Monarch Total RNA kit (New England Biolabs), and the High-Capacity cDNA Reverse Transcription kit (Thermo Fisher Scientific) was used for cDNA synthesis. Gene expression was measured with SYBR Green PCR Master Mix, 10 ng/µL cDNA, and respective primers listed in Supplement Table 1. Quantstudio 6 Flex Real-Time PCR System was used to run the experiment. ΔΔCT values were calculated by comparing the treated group with its respective untreated group.

### Protein lysates and Western blot

Preparation of protein lysates and Western blot analyses were performed as previously described [21]. Briefly, cell pellets were first washed with cold phosphate-buffered saline (PBS), lysed in RIPA buffer (Sigma) with protease inhibitor cocktail for 30 minutes on ice, and finally centrifuged ≥ 15,000g for 30 min at 4 °C to collect the supernatant as total soluble protein lysate. For muscle tissue, 30 mg of frozen tissue was first homogenized with 30–40 strokes in RIPA using a Dounce homogenizer (Wheaton Dounce Tissue Grinder, Catalog #357538), followed by shearing the tissue by passing through a 25-gauge syringe. For further lysis, the homogenate was rotated at 4 °C for ∼ 2 h, centrifuged at > 20,000g at 4 °C for 30 min, and the supernatant was collected as soluble muscle protein lysate. The lysates were quantified using BCA assay (Thermo Fisher Scientific). 20 µg of protein lysate was used for Western blot and Trans-Blot Turbo system (Bio-Rad Laboratories) was used to electro-transfer proteins to 0.45-µm PVDF membranes (Thermo Fisher Scientific) and immunoblotted with primary antibodies against GAPDH (1:2500, Abcam 9485), TDP-43 (1:2500, Abcam 190963), LC3B (1:2000, Abcam 192890), SQSTM1/p62 (1:15,000 Abcam 56416), p-TDP-43 (1:2500, Cosmo Bio USA: CAC-TIP-PTD-M01A), Ubiquitin (1:1000, Abcam 7780), and VCP (1:20,000 Abcam 11433 ). Secondary antibodies used were either anti-rabbit horseradish peroxidase (HRP) (1:5000) or goat anti-mouse HRP (1:5000).

### Immunohistochemistry

Immunohistochemical analyses were conducted on ASO-treated SMPCs and myotubes differentiated from SMPCs. SMPC were either cultured in matrigel coated 8-well chamber slides (Ibidi, IbiTreat 80826) for SMPC characterization or in 4-well chamber slides (IbiTreat 80426) for myotube differentiation study and ASO treatment on SMPCs. After the specified time, the cells were fixed in 4% paraformaldehyde (PFA). The fixed SMPCs were permeabilized with 0.2% Triton-X, blocked with Universal Block Buffer (UBB) followed by incubation at 4 °C overnight with primary antibodies for MF20 (1:100, DHSB), Pax7 (1:50, DSHB), SIX1 (1:1000, Novus Biologicals NPB2-52873), TDP-43 (1:200), LC3B (1:1000), SQSTM1/p62 (1:200), ubiquitin (1:100, Santa Cruz sc-8017), VCP (1:500), p-TDP-43 (1:2000), or galectin-3 (1:300, Santa Cruz A3A12). The cells were then washed three times with PBS and incubated with the secondary antibodies donkey anti-rabbit Cyanine Cy™3 (Jackson ImmunoResearch, 711 165 152); or donkey anti-mouse Alexa Fluor™ 568 (Thermo Fisher Scientific, A10042; 1:500) at room temperature for 1 h, then washed and mounted with DAPI-containing mounting media (Vectashield, Vector Laboratories, H-1200-10). All images were acquired with a Zeiss LSM 900 confocal microscope (Carl Zeiss). The mean fluorescence intensity was evaluated with ImageJ. Three images were acquired for each group, and at least 30 cells were analyzed.

### In vivo treatment of the VCP A232E mice with ASOs

Control non-targeting ASO and VCP ASOs were reconstituted in PBS to a final concentration of 10 mg/ml. VCP A232E mice were injected subcutaneously with control ASO, or VCP ASO at 50 mg/kg weekly. For the safety and tolerability experiment (Figure S1), 10-month-old VCP A232E were dosed with control ASO, or VCP ASO at 50 mg/kg weekly for 3 months and compared to WT mice. Mice were regularly monitored for weight loss, alertness, and activity. Weight loss of ≥ 15% compared to baseline was considered significant.

Serum was obtained from mice after 3 months of treatment with either control ASO, or VCP ASO, and used to measure alanine aminotransferase, aspartate aminotransferase, and creatine kinase. Organ weights (liver, kidneys, spleen) were collected and analyzed relative to body weight. For chronic treatment with VCP ASO to assess safety and efficacy, 6-month-old VCP A232E mice were injected weekly with control ASO or VCP ASO for 3 months and assessed for motor function. After 3 months of treatment, quadriceps muscle was collected for qPCR, Western blot, and histological assessment using H&E staining.

### Muscle Strength Measurement Rotarod accelerating speed test

The Rotarod test was performed using a five lane Rotarod system (Med Associates Inc.) to investigate motor coordination and learning skills by measuring the ability of the mouse to stay and run on the accelerated rod. Mice are trained to walk on a rotating rod and then assessed for their ability to maintain balance as the rod accelerates from 4 to 40 RPM for a maximum time of 5 minutes. The test is conducted over 3 consecutive days. Each day, the mice are assessed for three trials with a recovery phase of 1-2 min between trials. As soon as a mouse falls off the rod or starts to rotate with the Rotarod without running, the timer is stopped. The latency time to fall from the rod in seconds is recorded and average from 3 trials and the 3-day average is used for the analysis.

### Grip strength test

The grip strength test measures the muscle strength of the forelimbs in rodents. Grip strength is measured using a commercially available Grip Strength Meter (TSE Systems GmbH, Berlin, Germany). Each mouse is held by the base of the tail above a wire grid and gently lowered down until its front paws grasped the grid. The animal is then brought to an almost horizontal position and pulled back gently but steadily until the grip is released. Five trials testing with 1-minute intervals between trials are performed per animal. The force measurements are recorded from the five trials and averaged. The average of the three days of testing is used for the analysis [21].

### Inverted screen test

Inverted screen testing is used to assess the muscle strength of all limbs in mice. No prior training is required since the mouse has a natural motivation to hang on to the screen to avoid falling [3]. Each mouse is placed on the metal wire screen for ∼1 minute to allow it to adjust to the new environment. The mouse is then moved to the center of the wire screen. The screen is flipped quickly, but gently, so the mouse is hanging upside down. The time during which the mouse is suspended upside down on the wire screen before falling is measured. The maximum time for a single trial is set at 300 seconds. The mouse is allowed to rest for ∼1 minute before placing it on the wire again. The test is repeated for two additional trials, unless the mouse hangs on for 300 seconds (i.e., if the mouse hangs on for 210 seconds in trial 1 and 300 seconds in trial 2, then trial 3 was not performed). The hanging time in seconds is recorded, and the average of the 3 trials of testing is used for analysis.

### Hematoxylin and Eosin (H&E) staining

Quadricep muscles from mice are harvested, flash frozen in 2-methyl butane (Sigma) and embedded in optimal cutting temperature (OCT) compound. Tissue sections of 8 µm thickness are cut using a CryoStar NX50 (Epredia) microtome and subsequently stained with H&E stain following standard protocols. All images are acquired with BZ-X800 microscope (Keyence)

### Statistical analysis

Statistical analysis was performed using GraphPad Prism 9.1.2 software (GraphPad Software, Boston, MA). Data are presented as mean ± SEM. Multiple unpaired t tests were used for qPCR, WB densitometry quantification, and immunofluorescence image analyses of VCP R155H patient in comparison to WT iPSC-derived SMPCs. For WB analysis of WT and VCP R155H iPSC-derived SMPCs treated with and without VCP ASO and with Bafilomycin, two-way ANOVA followed by Šídák’s multiple comparisons test was used. One-way ANOVA test followed by Dunnett’s multiple comparisons test was used for qPCR and WB analyses in WT and VCP R155H iPSC-derived SMPCs treated with different doses of VCP ASO. Two-way Repeated Measures ANOVA followed by Dunnett’s multiple comparisons test was used for motor testing data analysis using Rotarod, grip strength, inverted screen tests, and for body weight analysis. One-way ANOVA test followed by Dunnett’s multiple comparisons test was used for qPCR and WB densitometry quantification, and organ weight analysis in treated A232E comparison to WT mice and unpaired t test was used for central nuclei quantification. For blood toxicology and liver enzyme analyses, one-way ANOVA followed by Fisher’s Least Significant Difference test was used. The level of statistical significance was set at *p = 0.05, **p = 0.01, ***p = 0.001, ****p = 0.0001.

## Acknowledgements

We thank the patients and families with VCP multisystem proteinopathy. We thank the Muscular Dystrophy Association for funding.

## Notes

### Competing Interest Statement

Michele Carrer, Paymaan Jafar-nejad, and Tamar Grossman are current or former paid employees of Ionis Pharmaceuticals.

