## Supplemental Figure and Table for "Antisense oligonucleotides targeting Valosin-containing protein improve muscle pathology and molecular defects in cell and mouse models of multisystem proteinopathy"

**Supplementary Material**

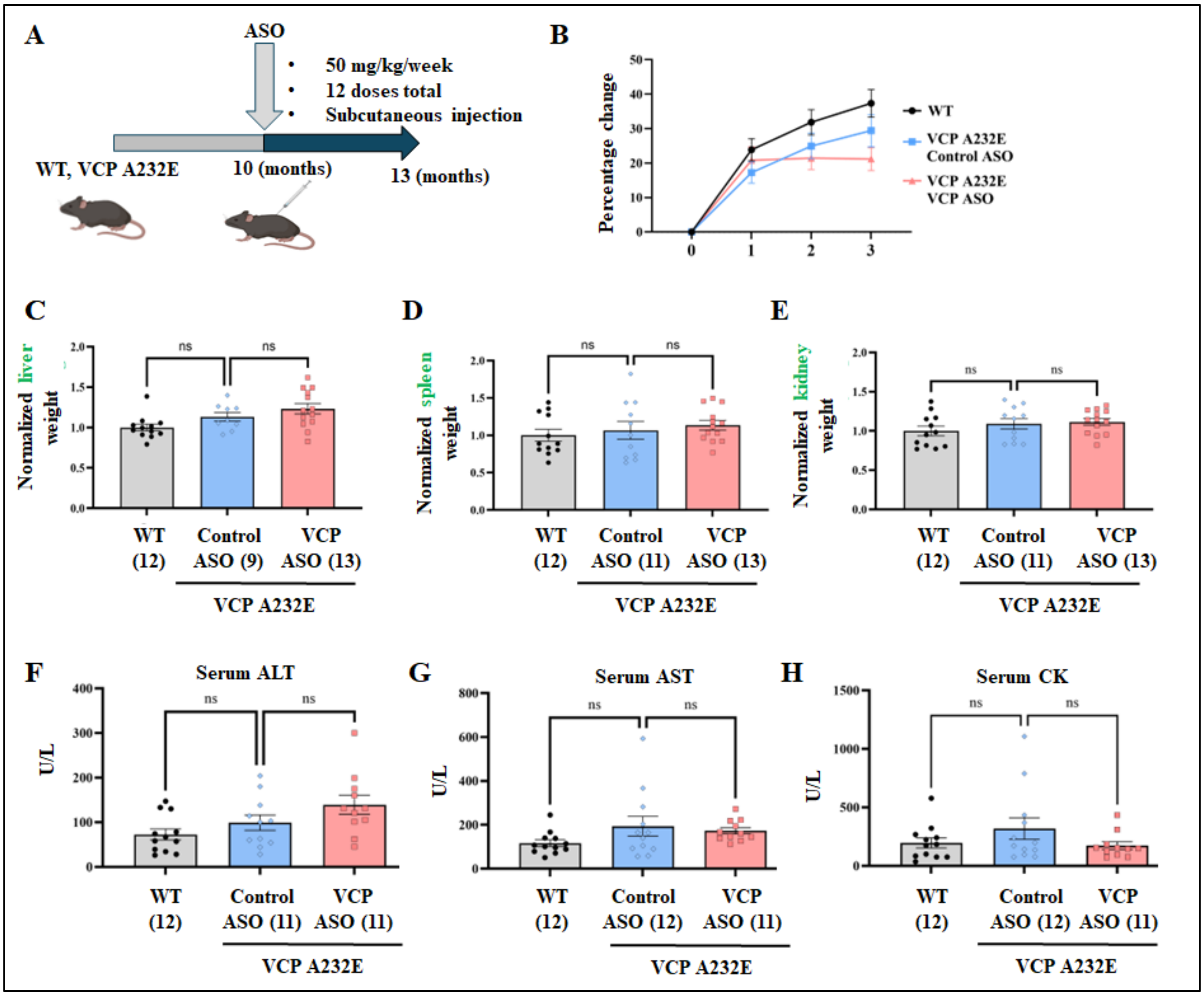


**Supplementary Figure S1. VCP ASO is well tolerated in VCP A232E mice at 50 mg/kg.** (A) Schematic illustration of the experimental design. 10-month-old VCP A232E mice (n=9-13) were injected weekly via subcutaneous injection with control ASO or VCP ASO at 50 mg/kg/week for a total of 12 weeks. (B) Body weight percent change for each treatment group. Measurements were recorded at baseline and monthly for 3 months. (C, D, E) The weight of key organs (liver, spleen, and kidneys) was measured at the end of the study and compared to the control ASO group. Data is presented as the mean ± SEM of the liver, spleen, and average of both kidneys normalized to mouse body weights for each treatment group. One-way ANOVA test followed by Dunnett's multiple comparisons test was used for statistical analysis (F, G, H). Mouse serum toxicology analysis: ALT: Alanine Transaminase, AST: Aspartate Aminotransferase, CK: Creatine Kinase, U/L: Unit/Liter. Data is presented as the mean ± Standard Error of the Mean (SEM).

**Supplementary Table 1: Sequences of gene-specific primers used for real-time PCR.**

| | | | |
|---|---|---|---|
| Species | Genes | Forward | Reverse |
| Human/ Mouse | GAPDH | 5′-CTCTGCTCCTCCTGTTCGAG | 5′-TGGTGTCTGAGCGATGTGG |
| Human | SQSTM1/p62 | 5′-TGTGTAGCGTCTGCGAGGGAAA | 5′-AGTGTCCGTGTTTCACCTTCCG |
| | LC3B | 5′-CAGCATCCAACCCAAAATCCC | 5′-GTTGACATGGTCAGGTACAAG |
| | TDP 43 | 5′-GATGGACGATGGTGTGACTGCA | 5′-AAGAACTCCCGCAGCTCATCCT |
| | VCP | 5′-GAGGAATCCTGCTTTACGGACC | 5′-GGCTTTACGAAGGTTGCTCTCAG |
| Mouse | SQSTM1/p62 | 5′-CTTACGGGTCCTTTTCCCAAC | 5′-TCCTCCTTGCCCAGAAGATAG |
| | LC3B | 5′-CGTCCTGGACAAGACCAAGT | 5′-CCATTCACCAGGAGGAAGAA |
| | TDP 43 | 5′- TTCATTCCCAAACCATTCAG | 5′-AGATGAACTGGATTACCACC |
| | VCP (WT) | 5′-CACAGTGGAAGGCATCACTGG | 5′-AAGGGCTGGGATCTGTCTCT |
| | VCP (A232E) | 5′-CAACGTGCTGGTTGTTGTG | 5′-AAGGACGATGCAAACAGCTT |
